# Off-target drift of the herbicide dicamba disrupts plant-pollinator interactions *via* novel pathways

**DOI:** 10.1101/2024.06.20.599889

**Authors:** Regina S Baucom, Veronica Iriart, Anah Soble, Matthew R. Armstrong, Tia-Lynn Ashman

## Abstract

Assessing the impact of herbicide drift on plant-pollinator interactions is crucial for providing insight into the causes of ongoing pollinator declines. The recent exponential increase in the use of the synthetic auxin herbicide dicamba, which is known to drift long distances following application, renders this concern especially acute. However, experimental data on the consequences of dicamba drift on plant-pollinator interactions are lacking from weed communities in natural settings. We assessed the indirect effects of dicamba drift on pollinator visits for 11 weeds of agricultural crops using a common garden field experiment, focusing on the potential for changes in pollinator abundance and alterations to both plant traits and patterns of pollinator visitation. We found variation among plant species in the extent of damage from dicamba drift exposure, and variation in how growth, flowering time, and flower displays were impacted, with some species showing negative impacts and others showing little effect. Pollinator frequencies were reduced in dicamba-exposed plots, and pollinator approaches and foraging visits were reduced for some weed species yet not others. Structural equation modeling revealed that the relationship between flower display and pollinator visits was disrupted in the presence of dicamba compared to control plots. Our study provides the most comprehensive picture to date of the impacts of dicamba drift on plant-pollinator interactions, with findings that highlight an underappreciated role of services supplied by weedy communities at the agroecological interface.

## Introduction

Weeds, defined as “plants that grow out of place” (1), have persisted in agricultural spaces since the beginning of agriculture (2), concomitantly providing crucial ecosystem services and leading to reduced crop yields (3, 4). Weedy plant communities that exist in the agroecological interface, or the non-managed areas surrounding agricultural fields, harbor beneficial insects, mycorrhizae, birds and other natural animal populations, and additionally reduce soil erosion and sequester carbon (5, 6). A particularly important benefit of these persistent weed communities are the resources they provide to pollinating insects throughout the growing season (7–9) and especially late in the season after crops have finished flowering (7). Diverse weed communities offer a variety of food resources that in turn benefit a broad range of pollinating insects (7, 10–12).

Regardless of their importance to pollinator health, weedy plants in the agroecosystem are most commonly viewed as pest species that should be rigorously managed (13). This management involves the use of herbicides designed to remove 90% of a given weed population (14). One of the consequences of herbicide use for weed control, however, is herbicide drift, where, during or post-application, the herbicide moves to nontarget areas surrounding crops, exposing the weed communities in the agroecological interface to low doses of herbicide (15, 16). The scientific focus on crop weeds and herbicides has long been on the management of resistance evolution rather than on the consequences of herbicide drift (17–19), meaning that we currently have a notable gap in our understanding of how sublethal doses of herbicide from drift influence the growth and flowering of weedy plant populations found within these ‘waste’ or unmanaged spaces on the crop edges (20, 21). Of particular importance is that we have an extremely limited understanding of how plant-pollinator interactions may be impacted following direct, low-dose herbicide exposures to the weed community (22). This knowledge gap is compounded by the decline of pollinating insects on the order of 50% in natural landscapes over the past 20 years (23), with intensive agricultural regimes (*i.e.,* the use of herbicide and large-scale farming) indicated as a key factor associated with this decline (23). Although dissecting the direct and indirect consequences of herbicide use and herbicide drift was identified as a key priority within the US’s pollinator research action plan (24), we still have a very limited understanding of the indirect consequences of herbicide application on pollinator health (25, 26).

Clarifying the role that herbicides may play in reducing the availability of floral resources to pollinating insects will continue to be of extreme importance as agriculture expands and changes to meet the needs of the global food supply. Agriculture currently accounts for 52% of the landmass in the US (27), and across this landscape, 73% of the pesticides applied yearly are herbicides (28). Recently, use of the phenoxy herbicide dicamba (3,6-dichloro-2-methoxybenzoic acid) for weed control has exponentially increased in the US given the adoption of dicamba-tolerant soy and cotton in 2016 (29, 30).

Unfortunately, dicamba, which mimics the natural plant hormone auxin, is known for its propensity to drift to nontarget areas (31), and in 2021 alone, there were 3,461 reports of dicamba drift from farmers and landowners (30). Although dicamba drift has the potential to negatively influence natural populations of plants and insects in agricultural landscapes, current studies of this dynamic revolve almost entirely around the impact of dicamba drift on crop yield (32–35), such that only a handful of projects have examined the potential that dicamba drift may alter the availability of crucial resources to pollinating insects (20, 36–38). Thus, the consequences of dicamba drift to natural plant and insect communities–in light of the recent shift in the US to the herbicide dicamba for weed control–is both an understudied and highly consequential issue in agriculture.

Currently, the available but limited work assessing the effects of dicamba exposure on natural, non-target plants demonstrates that low doses of the herbicide can delay flowering, reduce flowering duration, and reduce the total number of flowers produced (20, 36, 38). Strikingly, previous work has largely focused on the effect of dicamba drift using a single weed species (36), or by studying multiple weeds in greenhouse conditions (38). There are currently no examinations of the effect of dicamba drift on plant-pollinator interactions set in realistic field settings and using multiple ecologically relevant weed species. Thus, a major limitation of the work to date is that we do not understand how dicamba drift exposure alters plant traits across a number of relevant weeds, and how and if such trait changes may lead to cascading effects on plant-pollinator interactions. This has left a critical gap in our understanding of how the widespread shift to dicamba in agriculture may alter the availability of resources for pollinators.

To fill this knowledge gap, we examined plant-pollinator interactions in a weed community exposed to dicamba drift in field settings, focusing on the potential for changes in pollinator abundance and alterations to both plant traits and patterns of pollinator visitation. We simulated an off-target dicamba drift event at the flowering community level which was similar to what natural plant communities found at the agro-ecological interface experience. We originally sampled seeds from 11 weed species (Table 1) found in areas surrounding agricultural fields either in TN or KY (USA) (SI Appendix, Table S1) and planted replicates of each species into a common garden experiment, with half of the experimental plots exposed to dicamba drift (1% of the field dose) and the other half maintained as a control (Methods). We performed pollinator abundance screens and pollinator observations across experimental plots to determine whether dicamba drift altered the abundance of pollinating insects (categorized into 12 broad functional groups) or influenced how they approach flowers or forage for nectar or pollen. We likewise recorded plant traits such as size, leaf damage, flowering phenology, and flower display, and then used structural equation modeling to 1) examine the potential that dicamba drift indirectly altered floral visitation rates by directly influencing plant traits, and 2) determine if interrelationships between traits were altered by the presence of dicamba drift.

**Table 1.**
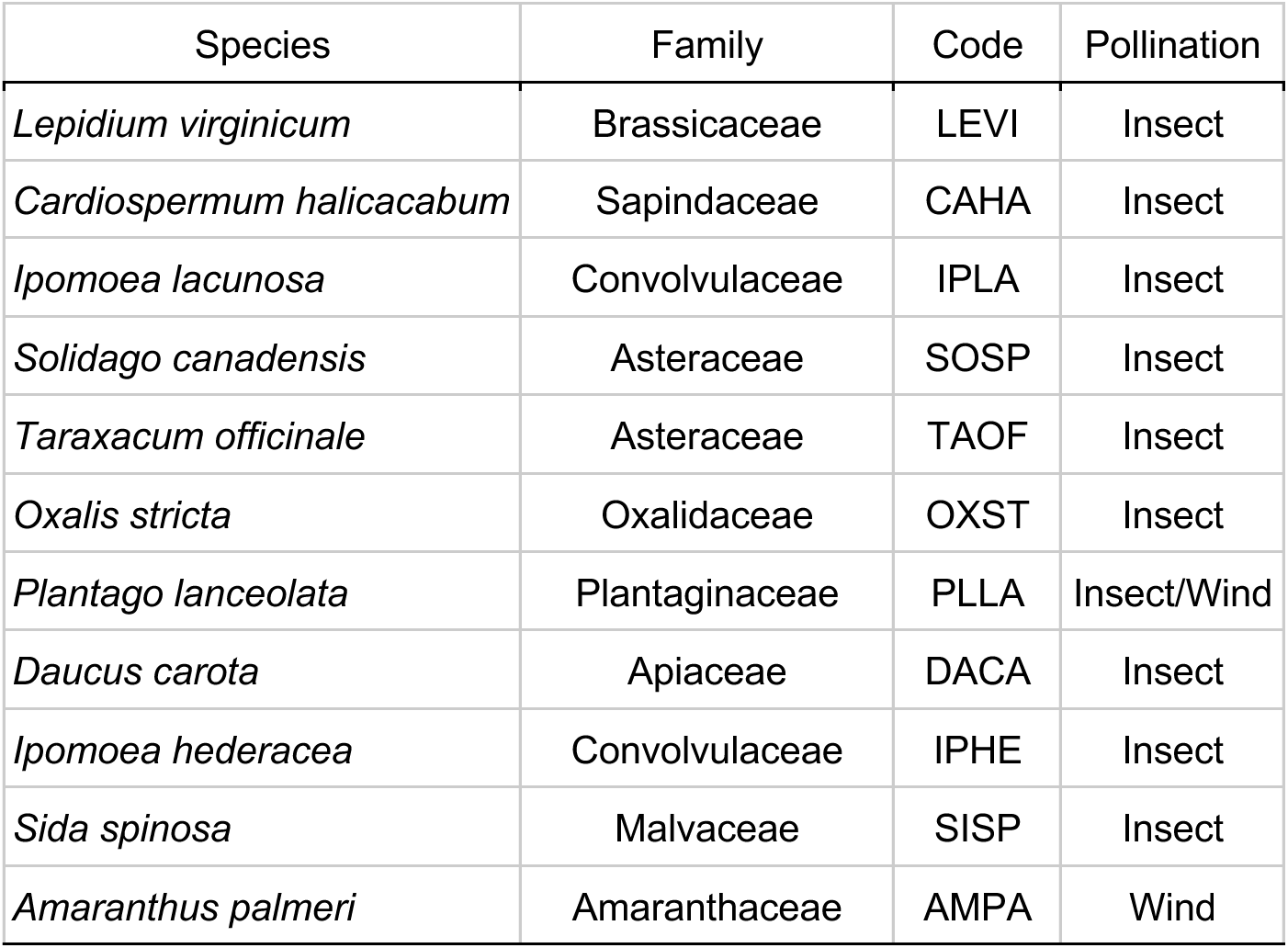
The eleven species used in this study to assess the impact of dicamba drift on plant traits and insect visitation rates. Each species, their family, and code used throughout the work is listed according to the United States Department of Agriculture PLANTS database. Mode of pollination (insect/wind) is also presented and was based on designations in (73), Hilty (http://illinoiswildflowers.info), and (74).

## Results

*Insect abundance*–We recorded a total of 161 pollinators across all control non-dicamba plots and 48 pollinators across all dicamba-exposed plots. This reduction in pollinator abundance in the dicamba exposed plots was significant (Figure 1*A*; treatment effect, *χ*^2^_1_ = 4.98, p = 0.03) with the general trend of reduced abundance across each of the functional groups considered (Fig 1*B*). Ants appeared to be the most affected functional group (Fig 1*B*), potentially due to altered relationships with *Daucus carota*. Removing ants from the analysis, however, did not qualitatively change overall results (treatment effect, *χ*^2^_1_ = 5.46, p = 0.02).

**Figure 1.**
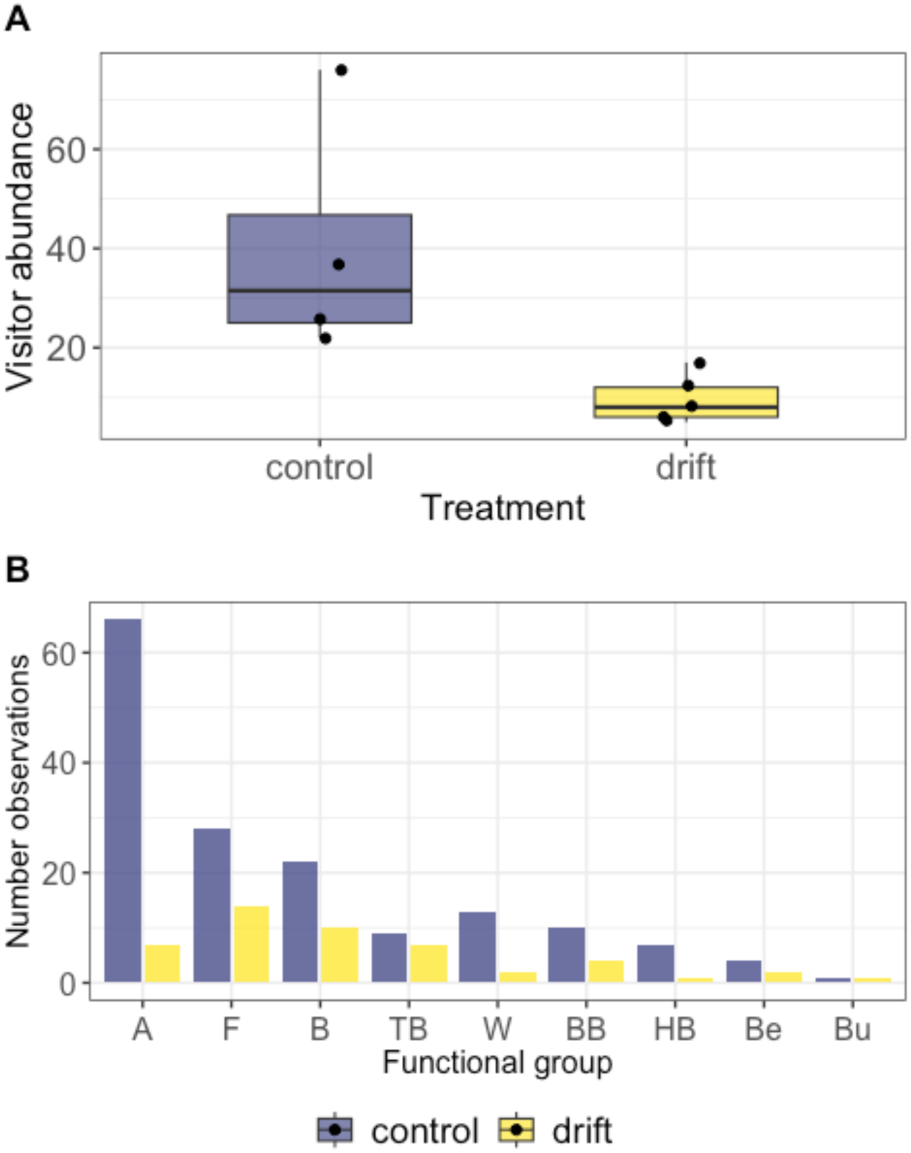
Pollinator abundance in dicamba drift exposed and non-exposed control plots. During 12 rounds of 20 minute surveys in each plot (total 240 mins/plot), the A) number of pollinating insects present was recorded, and each instance was placed into B) broad functional groups (A = ant, F = fly, B = Bee, TB = true bug, W = wasp, BB = bumblebee, HB = honeybee, Be = beetle, Bu = butterfly). Significantly more pollinators were present in control compared to dicamba-exposed plots (p = 0.03; Table S2).

*Pollinator approaches and foraging visits*–A total of 475 approaches and 10,072 foraging visits were recorded across surveys (see Fig S1 for overall visitation trends). The majority of approaches and visits occurred in the control plots (visits: 9456 control vs 616 dicamba; approaches: 310 control vs 165 dicamba). Flowers of *Daucus carota*, *Cardiospermum halicacabum*, and *Sida spinosa* were visited most frequently, experiencing a respective 72%, 23% and 3% of the overall visits (see Fig S2 for visits per species across surveys).

When pollinator approaches were modeled as a binary response we found a significant treatment by plant species interaction (*χ*^2^_7_ = 20.46, p = 0.004; Table S2). Three species were responsible for this effect, with *Sida spinosa, Ipomoea hederacea,* and *Ipomoea lacunosa* in the dicamba treated plots being 58%, 54%, and 28% less likely, respectively, to be approached by a pollinator compared to a plant of these species in the control plots (Fig 2*A*). Whether or not a plant was visited likewise showed a significant treatment by species interaction (*χ*^2^_7_ = 25.11, p < 0.001), but this effect was driven entirely by *D. carota*, which had 0 visits in the dicamba present environment (species x treatment interaction without *D. carota*: *χ*^2^_6_ = 8.45, p = 0.21; Table S2). When pollinator approaches and visits were modeled as rates, we found that approach rates were not influenced by dicamba exposure (treatment effect, *χ*^2^_1_ = 0.02, p = 0.88; Table S2) nor did approach rates differ across species due to the presence of dicamba (species x treatment effect, *χ*^2^_1_ = 7.10, p = 0.52; Fig 2*B*, Table S2). However, we found a significant species by treatment interaction for visit rate (*χ*^2^_1_ = 31.38, p < 0.001), driven largely by lower visit rates for *C. halicacabum*. There was no evidence of an overall treatment effect for visit rate (*χ*^2^_1_ = 0.01, p = 0.94).

**Figure 2.**
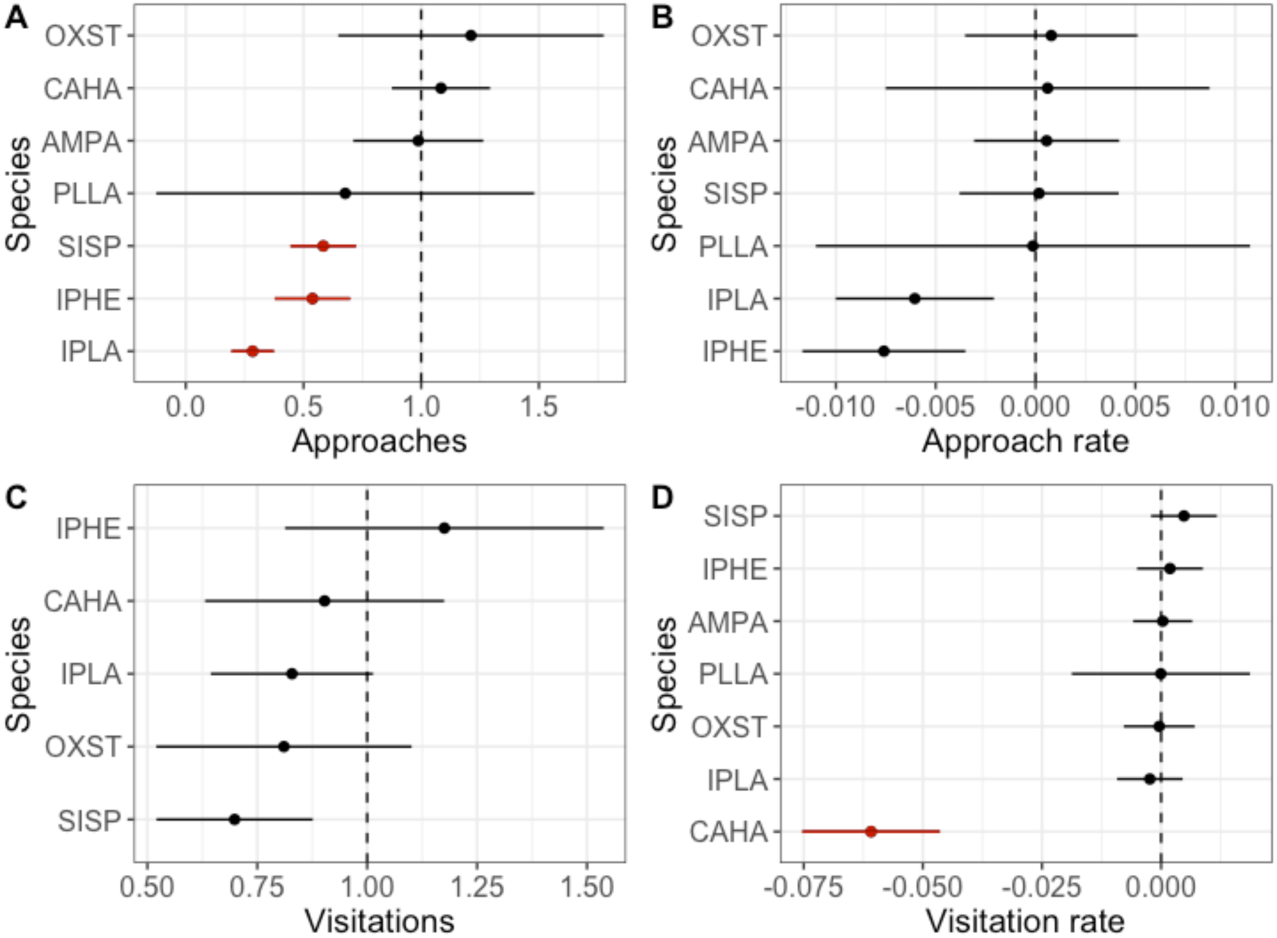
The results of twelve survey rounds recording floral approaches (an approach but no foraging) and visitations (foraging within the flower). Approaches and visitations were modeled both as a binary response variable (yes/no) for each plant across each of the survey rounds (A, C), and likewise as a rate (number of approaches or visitations per flower per hour (B, D)). In A and C, the risk ratio (ratio of probabilities) for insect approaches or visitations are shown, respectively, where values below 1 indicate that insects have a lower probability of approaching or visiting a flower in the dicamba compared to the control environment. In B and D, contrast estimates (±SE) depict the difference between dicamba exposed and control plants for each species, with statistically significant differences shown in red. Species are designated by their four-letter code as in Table 1. Approach and visit rates shown in B and D were first back-transformed prior to plotting. Approach and visitation contrasts are not shown for DACA (*Daucus carota*) given that we recorded no floral approaches or visitations in the dicamba present environment, and many approaches and visitations in the control environment (see *Results*). We likewise do not graphically depict PLLA or AMPA in C given that few visitations in the dicamba environment occurred for these species, leading to large standard errors.

*Plant traits*–Dicamba exposure led to both smaller plant size and significant leaf damage across species (size at three weeks, treatment effect, F_1,7.3_ = 7.71, p < 0.03; leaf damage, treatment effect, F_1,6.3_ = 152.28, p < 0.001; Table S3) with a significant species by treatment interaction for both traits indicating that some species were more impacted by dicamba drift than others under field settings (Table S3).

*Amaranthus palmeri*, *Sida spinosa, Ipomoea hederacea* and *I. lacunosa* exhibited the greatest reduction in growth compared to plants grown in control plots (Fig 3). All species displayed significant leaf damage, ranging from 5-15% damaged leaves across species, with *Cardiospermum halicacabum, I. hederacea* and *I. lacunosa* exhibiting the most damage (Fig 3).

**Figure 3.**
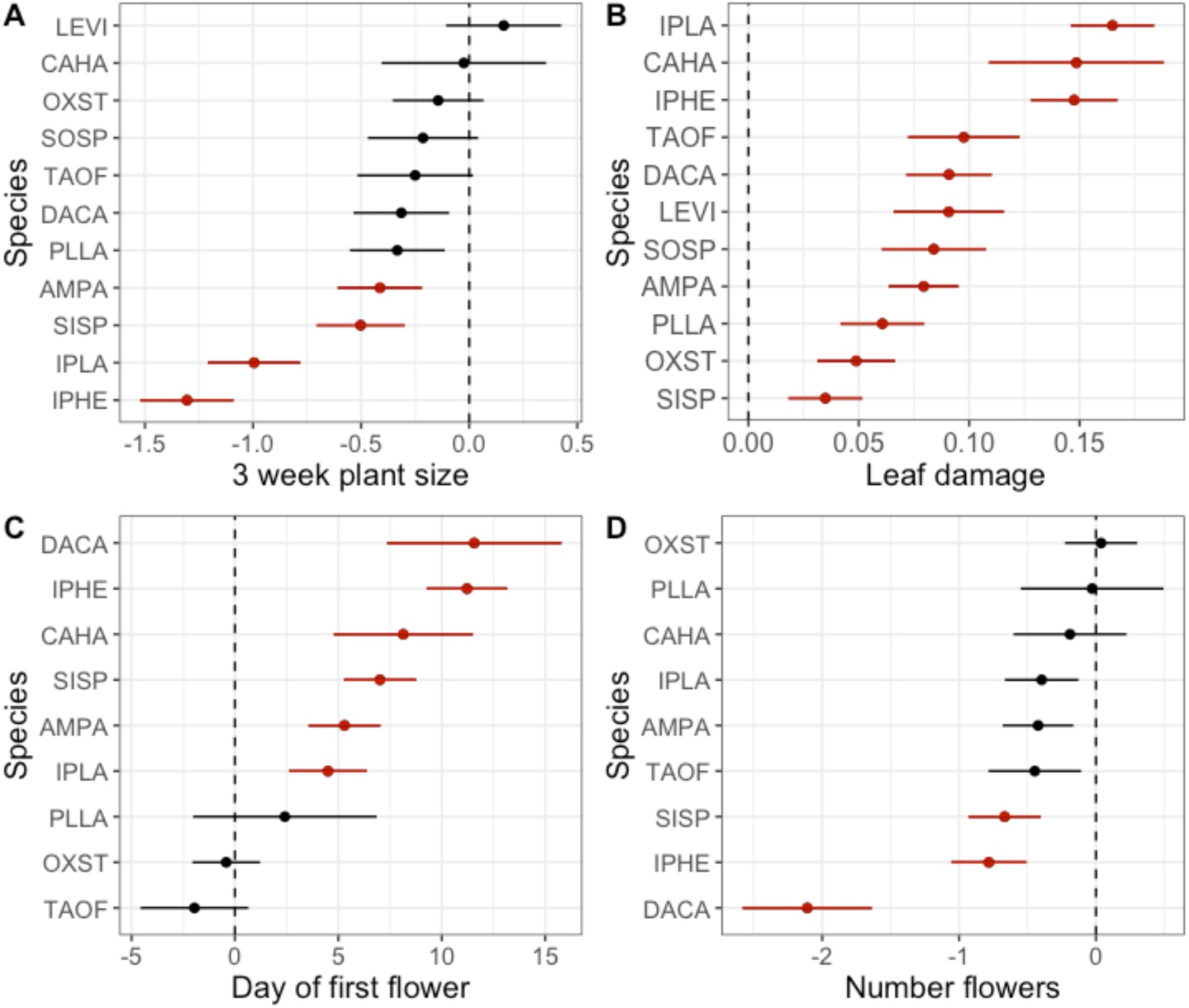
Plant species vary for A) size at three weeks post-dicamba exposure, B) the amount of leaf damage exhibited, C) the first day of flowering, and D) the total number of flowers produced over the experiment. Contrast estimates (±SE) show the difference between dicamba exposed and control plants of each species, with statistically significant differences depicted in red. The treatment by species interaction was significant for each of the traits (A-D) with p < 0.001 (Table S3). Species are designated by their four-letter code as in Table 1. Estimates were first back-transformed prior to plotting.

Flowering was significantly delayed in the dicamba exposed plots (treatment effect, F_1,10.3_ = 17.55, p = 0.002). Nine of the 11 species flowered in either the control or dicamba plots during the experiment, with six of the nine species exhibiting significantly delayed flowering in the presence of dicamba (avg days the onset of flowering was delayed: *Daucus carota* (11.6)*, Ipomoea hederacea* (11.2)*, Cardiospermum halicacabum* (8.1)*, Sida spinosa* (7)*, Amaranthus palmeri* (5.3), and *I. lacunosa* (4.5); species x treatment, F_1,6.3_ = 4.84, p < 0.001; Fig 3). Dicamba drift exposure also led to a significant reduction in the total number of flowers produced over the experiment (treatment effect, F_1,7.9_ = 6.02, p = 0.04; Table S3) with *S. spinosa, I. hederacea,* and *D. carota* exhibiting significantly fewer flowers in the presence of dicamba and *Ipomoea lacunosa, Taraxacum officinale,* and *Amaranthus palmeri* showing a non-significant reduction in flower number in the presence of dicamba (species x treatment, F_1,207_ = 3.76, p < 0.001; Table S3, see Fig S2 for trends in flowering for each species in each environment across survey rounds). Preliminary analyses evaluating phylogenetically controlled models for each plant trait (as well as pollinator approaches and visits) exhibited higher AIC values than analyses that were not phylogenetically corrected, thus we report non-phylogenetically corrected models in the main text (see *SI Appendix*, Methods S1; Fig S3, Table S4).

*Structural equation modeling–*We examined the hypothesis that dicamba exposure indirectly altered pollinator visit and approach rates through direct effects on plant traits using structural equation modeling. We developed three hypothesized models (Models A-C, Fig S4) to identify which of the plant traits were most influential following dicamba exposure. None of the three hypothesized models differed significantly from the observed data (Table S5), nor did the models vary significantly from one another (p > 0.05 for each model comparison). Model B showed a lower AIC value than the full model (Model A), indicating both a better fit to the data and that dicamba did not directly influence the first day of flowering or total number of flowers, as these two pathways were present in the full model but not in Model B. We present results using Model C, a reduced form of Model B (Methods; SI Appendix, Fig S4, Table S5), since it exhibited the lowest AIC value of the three and was the simplest model with the fewest parameters (Fig 4*A*).

**Figure 4.**
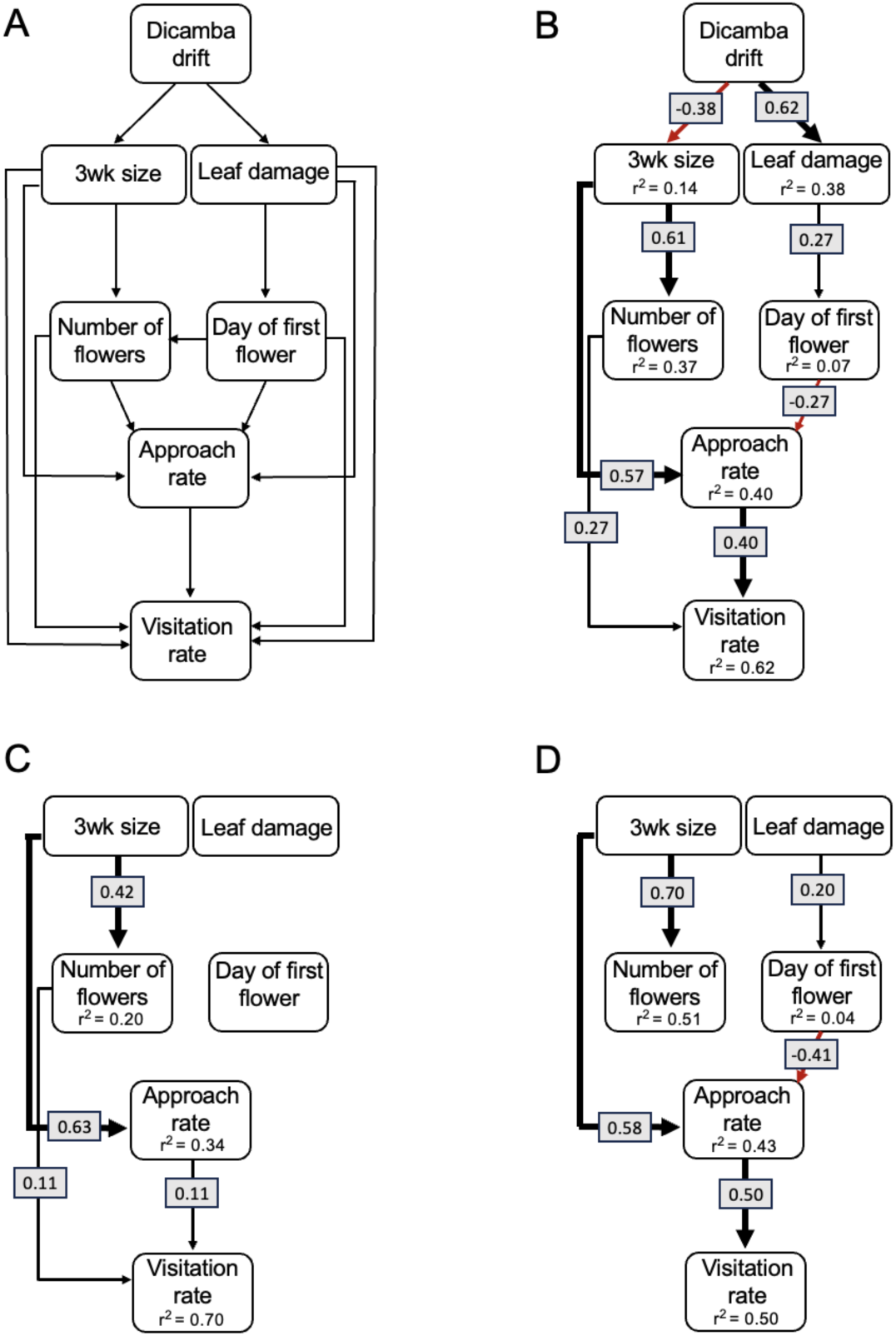
Structural equation modeling identified the indirect effect of dicamba on pollinator visit rate following dicamba drift exposure (dicamba treatment). In A, the hypothesized structural equation model (Model C, Fig S4) depicting potential relationships between dicamba exposure, plant traits, and pollinator approach and visit rates is shown; each hypothesized relationship is indicated by the direction of each arrow. This model provided a fit to the data as determined by d-Sep tests in piecewiseSEM (see Fig S4 and Table S5 for comparison of hypothesized models). Results of the overall model are shown in B, with significant (p < 0.05) standardized path coefficients for each relationship shown. Red lines indicate negative relationships whereas black lines depict positive relationships between either the exogenous variable ‘dicamba drift’ with endogenous variables or relationships between endogenous variables themselves. The standardized path coefficients shown here depict direct relationships between traits, and direct, indirect, and total effects for each trait are presented in Table S6. The same general model was used to evaluate the strength of the relationships between traits within the control environment (C) or within the dicamba environment (D). Model fit information is presented in Table S5 for each model.

Overall, Model C revealed three separate pathways by which dicamba exposure indirectly affected pollinator visit rates (Fig 4*A*; Table S6) with a weak but negative total effect of -0.17 (sum of the indirect and direct effects across pathways, Table S6). The first pathway indirectly influenced visit rate by -0.02 (Table S6; Fig 4*B*) and was mediated through leaf damage, day of flowering, and approach rate. The other two pathways were mediated through dicamba’s effect on plant size and contributed the majority of the negative indirect effect (-0.15, Table S6; Fig 4*B*). The approach rate and number of flowers both directly influenced the visit rate, with approaches and flowers positively influencing visits (0.40, 0.27), whereas the effects of plant size, day of flowering and leaf damage were indirect and were mediated through their effects on approach rates (Fig 4*B*).

We further found that relationships between traits differed between environments. When performing the SEM using only plants from the control environment, we found a moderate but significant positive relationship between the number of flowers and visitation rate (0.11, Fig 4*C*). This relationship was not present in the dicamba environment, where leaf damage directly positively influenced the day of flowering; plants with more damage flowered later (0.20; Fig 4*D*). Additionally, there was a negative relationship between flowering day and approach rate in the presence of dicamba (-0.41; Fig 4*D*) such that plants exhibiting earlier flowering exhibited higher approach rates. A few central relationships between traits remained the same between environments: in both the control and dicamba environment, three week size positively influenced both the number of flowers produced (0.42, 0.70; control vs dicamba environment, Fig 4*C-D*) and the approach rate (0.63, 0.58; control vs dicamba environment, Fig 4*C-D*). The approach rate in both environments significantly and positively influenced visitation rates (0.11, 0.50; control vs dicamba environment, Fig 4*C-D*).

Given the absence of a relationship between number of flowers and pollinator visits in the presence of dicamba compared to control plants, we elected to examine this relationship for each species separately to determine if a particular species was driving this change. To do so, we performed simple linear regressions between pollinator visits and number of flowers in both the dicamba present environment and the control environment for each species separately (*data not shown*). We found a significant positive relationship between visits and number of flowers in the control but not dicamba present environment only for *Cardiospermum halicacabum* (Fig 5*A*), indicating that the altered relationship was largely due to dicamba’s impact on the interactions between this species and its pollinating insects. The change in pollinator visits is likely attributable to a marked decline in visitations by bumblebees and wasps in the presence of dicamba (Fig 5*B-C*).

**Figure 5.**
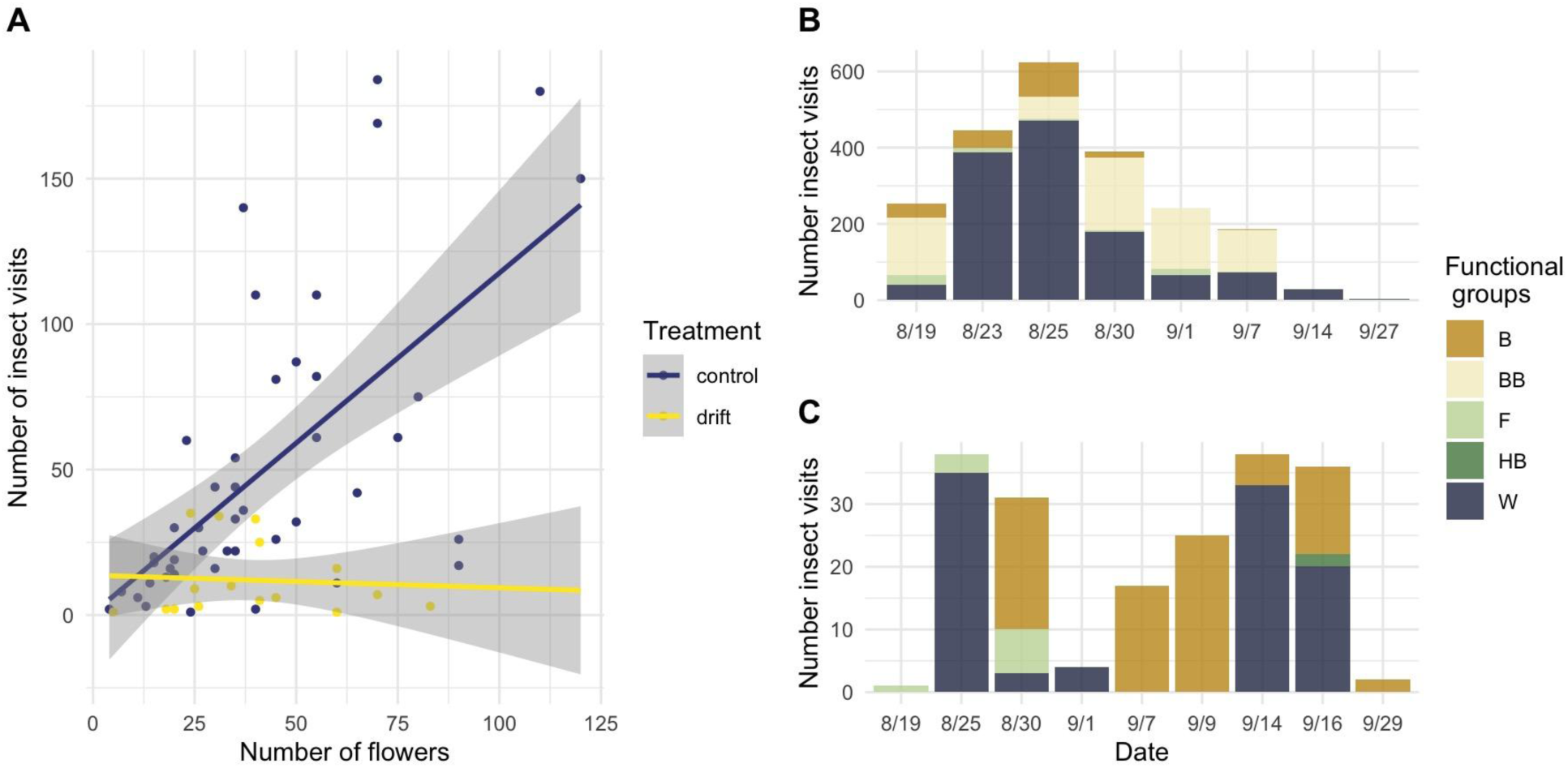
The relationship between (A) number of pollinator foraging visits and number of flowers in *Cardiospermum halicacabum* and the number of pollinator foraging visits per functional group visiting this species over the course of the experiment in the (B) control and (C) dicamba exposed plots. The relationship between the number of flowers and number of visits was significant in the control environment (F_1_ = 28.19, p < 0.001) but not the dicamba present environment (F_1_ = 0.39, p = 0.53).

## Discussion

Our analyses of plant traits of 11 common agricultural weed species exposed to dicamba at low doses, combined with pollinator abundance screens and an assessment of plant-pollinator visits provides the most comprehensive picture to date of how dicamba drift alters the floral rewards that weeds at the agroecological interface provide and how the the pollinator community responds to these changes. We show reduced pollinator abundance in experimental plots treated with dicamba drift – an effect that was largely present across all pollinator functional groups – and further found that while all weed species showed signs of leaf damage following dicamba exposure, species varied in whether this damage led to cascading effects on growth, flowering, and ultimately pollinator approaches and visitation rate. Our analysis further demonstrated that plant-pollinator interactions are altered primarily due to dicamba’s effects on flower productivity, and importantly, the oft-reported relationship between floral display and pollinator visits was disrupted in the presence of dicamba for at least one species. Overall, we document variability in responses to dicamba drift across weed species, and this result, in concert with evidence of reduced pollinator abundance, suggests that there are multiple ways in which dicamba drift can disrupt plant-pollinator interactions and that the dynamics of this disruption vary according to weed species.

### Dicamba drift reduces pollinator abundance and alters pollinator interactions

Our pollinator abundance surveys found a 70% reduction in pollinating insects within dicamba-exposed experimental plots compared to the control plots. Because we controlled for plot-level floral display in our analyses, the reduction in pollinators likely stems from overall floral compositional changes caused by differences in flower production among species. For example, while most weed species exhibited reductions in flower number (*e.g.*, 20-96% reduction in flowers produced in the dicamba exposed plots; SI Appendix, Fig S1, Fig S2), two species produced *more* flowers in the presence of dicamba (*e.g*., *Oxalis stricta* and *Amaranthus palmeri* produced 1.5 and >2 times the number of flowers in the dicamba exposed plots, respectively, albeit not significantly so).

Further, we found reductions in the likelihood pollinators would approach (*i.e.* a non-foraging visit) flowers of dicamba-drifted plants for a handful of species (*Sida spinosa*, *Ipomoea hederacea*, and *Ipomoea lacunosa*), but did not find differences in the approach rate–a metric that is standardized by total flower number per plant–across any species. The lowered likelihood of approaches of *S. spinosa*, *I. hederacea*, and *I. lacunosa*, but lack of changes to approach rate is likely due to dicamba-induced alterations in floral display, since each of these three species flowered later and exhibited fewer flowers in the presence of dicamba. Together, these results suggest that reductions in floral display from dicamba exposure cause a reduction in floral approaches for some weed species but not others. Intriguingly, when examining foraging visits (*i.e.* a pollinator approaches and then forages for pollen or nectar), we found that all species were equally likely to be visited between treatments, but that the rate of visitation was significantly reduced for one species (*Cardiospermum halicacabum*). As with approach rate, visitation rate is standardized by the total number of flowers each individual produced across surveys, potentially suggesting that the rate of visitation is reduced in *C. halicacabum* not because of changes in floral display, but instead due to reductions in the quality of the nectar or pollen or perhaps changes in the volatile organic carbons (VOCs) that are released by plants and attract pollinators (39).

Our work both expands and adds nuance to prior knowledge on dicamba-induced alterations to plant-pollinator interactions. Egan et al (20) report reductions in some arthropods (aphids and potato leaf hoppers) in crop edge communities exposed to dicamba drift, but found no changes in the abundance of insects considered potential pollinators (*i.e.*, small Hymenoptera species). However, Egan et al (20) used a modified leaf blower to vacuum insects from within plots, appearing to capture small, non-flying arthropods such as ants, leafhoppers, beetles, and crickets, whereas we performed visual surveys of pollinating insects present anywhere within the experimental plot (including flying near or on our focal plant species). Further, Bohnenblust et al (36) assessed pollinator visitation in a single weed species (*Eupatorium perfoliatum*) exposed to dicamba drift and reported significant reductions in pollinator visits in dicamba-drifted plants; however, this work was performed on potted plants arranged in an array and supplemented with a honey bee hive, meaning that the reduction in pollinator visits reported in this context might not be generalizable to agricultural field settings in which multiple flowering species are present. Altogether, our results, from a replicated field experiment using 11 weed species, and taking advantage of the natural pollinator community, confirms predictions from the work of Iriart et al (2022) by showing that decreased attractiveness of some species following dicamba exposure was responsible for fewer pollinator approaches, and that potential changes in the quality of floral rewards may underlie the reduced visitation rate for one species. Thus, our work highlights that dicamba at drift levels can significantly disrupt plant-pollinator interactions for some weed species, but that generalizations about the extent of the impact from one weed to another cannot easily be drawn.

### Dicamba drift impacts plant-pollinator interactions through effects on plant growth and flowering

Our structural equation model demonstrated that the relationship between flower number and pollinator visits is disrupted by the presence of dicamba. In plant communities, larger floral displays act as visual cues that attract pollinators to an area, thus increasing the number of visits (40–42). That this relationship was disrupted in the presence of dicamba demonstrates that floral cues, which have evolved through millennia of plant-pollinator interactions, can be altered given a single exposure of dicamba drift early in the plant life cycle (*i.e*., prior to flowering). We also demonstrated a generalized mechanism by which dicamba indirectly alters plant visitation rate–the herbicide led to reduced plant size, which consequently reduced flower production and, at least for one species, pollinator visit rates. That herbicide exposure more generally decreases flower production in crop weeds is not in itself a novel finding (43, 44), but the demonstration of a link between reduced flower number and pollinator visits as a consequence of dicamba exposure, along with an understanding of which traits influence this relationship, is novel.

Additionally, we uncovered a negative correlation between the day of flowering and approach rate in the presence of dicamba, but not in the control treatment. We hypothesize that this relationship could be due to a dicamba-induced mismatch between the phenology of the plants and the pollinators. Specifically, plants in the dicamba treatment flowered between 5-15 days later (on average) than control plants, such that earlier flowering individuals in the dicamba environment were likely flowering when the pollinator community was nearing its peak, whereas later-flowering plants potentially experienced lower approach rates due to declining activity of the pollinator community. Two species, *Ipomoea hederacea* and *Sida spinosa* were the most likely contributors to this pattern, as both exhibited significantly later flowering in the presence of dicamba and relatively high approach rates when flowering commenced (data not shown). Interestingly, our structural equation model demonstrated a positive relationship between the amount of leaf damage a plant exhibited and flowering phenology. Thus, plants that experienced significant leaf damage following dicamba exposure also exhibited later flowering, which then led to a reduction in pollinator approaches. Our results confirm past work showing herbicide-induced delays in flowering and predicted shifts in flowering community (38, 43–46), yet significantly expands our understanding of how this delay in flowering can influence pollinator pool foraging behavior.

### Implications for the conservation of plant-pollinator interactions in agroecological spaces

Pesticide use in agriculture–along with parasites, disease, global climate change and the reduction of natural habitat–is one of the key contributors implicated in pollinator declines (47). Our results highlight that dicamba, which has no currently known direct effect on pollinating insects (48, 49), indirectly influences pollinator health by reducing the abundance and complexity of floral resources that beneficial natural habitats surrounding crops provide to pollinators. These results offer several insights that can inform pollinator conservation efforts. First, we show that dicamba drift influences plant-pollinator interactions given its effect on plants across two main axes – reductions in flower number and delays in flowering time. However, we also show variation in flowering phenology and flower production across the species we examined. This suggests that practitioners planting buffer vegetation around crops to supplement pollinator diets (*e.g.* prairie strips (50) and similar mitigations) should place an emphasis on using diverse and varied plant communities to ensure the pollinator community has access to food resources given that dicamba impacts weed species unequally, and given that variation in pollen diets are associated with longer bee lifespan (10, 11) and increased resilience against pathogens (12, 13). Further, beyond visual cues, our data also suggest that aspects of plant quality can be impacted by dicamba, given that the visit rate (which controlled for total number of flowers present) was lower for one species following dicamba drift. Therefore, an important consideration when optimizing plant communities at the agroecological interface is understanding how dicamba drift may influence the quantity and quality of floral resources (*e.g.* pollen and nectar content and composition)–an area almost completely lacking in knowledge (but see (36)). Finally, in addition to the immediate ecological consequences (*i.e.* reduction and delay of resources), the alterations in plant traits has the potential to lead to new opportunities for selection and hence influence evolutionary changes. For example, pollinators can impose selection on both floral display and floral phenology; the plastic shifts that we demonstrate in these traits may impose natural selection in opposition to that from pollinators (51). Thus, practitioners and conservationists aiming to develop robust and stable plant communities at the agroecological interface should likewise study the potential for unexpected evolutionary changes given ecological consequences of herbicide drift (52).

## Methods

### Study species

We collected seeds from 11 wild flowering plant species (9 plant families) found within weed communities at the edge of soybean, maize, or fallow fields located in central Tennessee or western Kentucky during 2018 and 2019 (Table 1). Dicamba use in both areas of TN and KY began in 2017 and was predicted to increase by local extension agents (personal communication, Baucom). Mature seeds were collected from 1-3 populations per species (see SI Appendix, Table S1 for location and population information). Most species are insect-pollinated, with one wind-pollinated species, the latter of which pollinators will visit if resources are limited (Saunders 2018; personal observation, Ashman).

### Field experiments

On June 1, 2021 we planted six replicate seeds of each species in 11 tilled plots (6 x 11 x 11 = 726 seeds total) at the Matthaei Botanical Gardens (MBGNA) in Ann Arbor, MI. The plots were each at least 50 m away from one another and each were embedded within prairie landscape or grassland. Prior to planting, we bulked seeds of each species across populations thereby combining the sources and seeds were randomly allocated to plots from this bulk source. Replicates of each species were also planted in the greenhouse at this time, watered daily, and kept in standard conditions of 75F and 12 hr of artificial light. Since germination in the field was low, we replaced non germinating seed with transplants of each species on June 22-23 to ensure complete 6 individual plants per species per plot.

We applied 1% of the field dose of dicamba (3,6-dichloro-2-methoxybenzoic acid, Albaugh, LLC, Ankeny, IA) to plants within five randomly chosen plots and a 0% control to another four plots four weeks post transplantation on July 21, 2021. The 1% dicamba treatment represents a particle drift rate (*i.e.*, when herbicide particles travel by wind (53). We applied the field dose to the final two plots (561 g active ingredient per hectare) to ensure the efficacy of the dicamba formulation used. We included ‘Preference’ surfactant (non-ionic surfactant blend, WinField Solutions, St. Paul, MN) in all treatments (dicamba or control) at 0.1% v/v. Plants were sprayed from one pass until just wet using a handheld multi-purpose sprayer set to a medium-fine mist with an operating pressure of 40-45 PSI. We used a handheld multipurpose sprayer rather than a CO_2_ sprayer, which maintains a constant application rate, to simulate the variation that would occur in light of an off-target drift event in field conditions.

We performed twelve rounds of pollinator observations in each plot during which we recorded both approaches and visitations to flowers and abundance in the plot. Pollinators were grouped according to 12 broad functional groups (54). These include known pollinators plus other insects that visit flowers but may or may not pollinate flowers (*e.g.* ants (55)). During a round of pollinator observations, at each plot, we counted the number of flowers present on each individual experimental plant (target individual) for a measure of floral display. We then began recording visitations by visually scanning the plot and recording any visitations (forages within the flower) or approaches (coming in close proximity to a flower but not landing or completing a true visitation). Approaches capture the degree to which insects are attracted to a flower, whereas visits capture insect foraging. Observations were performed for 20 min/plot. Plants with open flowers but no visitations or approaches were scored as ‘0’ during each observation round. We performed one round of observations across all plots twice per week, and randomized the time of observation across plots during the twelve rounds (observations occurred within a window between 8am and 1pm each day).

Pollinator abundance surveys were likewise performed during 12 separate occasions (1 round/day, 20 min/plot) by recording the number and type of each pollinator present anywhere within the plot, again according to 12 broad functional groups. During these abundance surveys, at each plot, we counted the number of flowers present on each individual experimental plant (target individual) for a measure of floral display, and we likewise counted flowers of any non-experimental plant within the plot (labeled as ‘background’) as well as individual, non-experimental plants immediately surrounding the plot (labeled as ‘vicinity’).

Immediately prior to applying the treatments, we recorded the total number of leaves on each plant and the width of the largest leaf (cm) and used the product of these two traits to estimate pre-treatment plant size as in (38). Three weeks following the treatment application we again recorded the total number of leaves and the width of the largest leaf and used the product of these to estimate post-dicamba plant size. We additionally recorded the number of leaves showing signs of dicamba damage, typically leaf ‘cupping’ where leaves are wrinkled and take on a cupped shape (56). We likewise recorded the presence and survival of each individual throughout the experiment. Plants within a few plots experienced vole herbivory which was recorded. To determine whether dicamba treatment influences flowering phenology and floral display within a weed community, we recorded the day of first flowering and the number of flowers present on each plant three times per week from July 23 to September 29.

### Data analysis

We performed all analyses in R version 4.3.3 (57), RStudio (58), and used the ggplot2 package to create all figures (59). For each of the plant traits, we used mixed-effects linear models from the lme4 package (60), and modeled data from the control and drift treatments only. We first used the bestNormalize package (61) to examine all response variables for normality and to determine the most appropriate transformation to meet model assumptions. We tested the significance of fixed effects with Type III sums of squares using the Anova function (car package, (62)) and estimated denominator degrees of freedom using the Satterthwaite method (63).

To determine the effect of dicamba drift on each of the plant traits – plant size three weeks post treatment, leaf damage, first day of flowering, and total number of open flowers – we analyzed the model as:

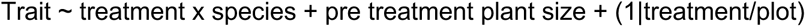

with treatment, and species included as fixed effects and plot included as a random effect. Pre-treatment size was included as a covariate in each model. We included the random effect of ‘transplanted’ preliminarily to account for any effects of transplanting individuals from the greenhouse but did not retain this term in final models since no effect of transplantation was uncovered. We used contrast estimates to classify species-specific patterns in each of the traits between treatments (*e.g.*, growth in drift subtracted by that in control) and identified estimates that were significantly different from zero using the contrast function (64).

We used a generalized linear mixed model with a poisson distribution to examine the potential that dicamba drift altered pollinator abundance. The dependent variable was the number of insects summed across survey rounds per plot (floral visitor ‘frequency’), and insect functional group was used as the independent variable in the following model:

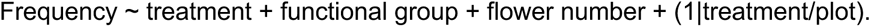

A ‘plot level’ flower number (flower number summed across plants and rounds) was used as a covariate to control for differences in floral display among plots, and we examined results using the flower number of all plants (target, background, and vicinity) or just the target plants. The results were qualitatively the same, and so we report results using total flower number from all flowering plants as the covariate. Plot was considered a random effect in the model and nested within treatment, and treatment, functional group (*e.g.*, ant, bee, etc), and flower number were considered fixed effects.

To examine the influence of dicamba drift on pollinator approaches and visits to flowers, rates, we initially used generalized linear mixed models (glmer) with Poisson distributions, but discovered that data were overdispersed and negatively skewed given a high proportion of zeros in the dataset (∼60%). Therefore, we performed analyses of approaches and visitations in two ways: first, we modeled approach and visitation as a binary ‘yes/no’ response variable for each plant across observation rounds with a binomial error distribution, and second, we modeled approach and visitation rates (number of approaches or visits per flower of the plant visited per hour) with a quasi-Poisson error distribution. In the analysis of binary response data, we included flower number of each plant as a covariate to control for differences in floral display among species. Given a lack of model convergence when the plot effect was included as a random variable, we used generalized linear models (glm) in base R for these analyses, with plot nested within treatment and the interaction between species and treatment considered as fixed effects. The following general model for both types of analyses was used:

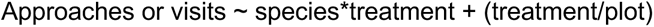

In each analysis, the treatment by species interaction was of interest since that would indicate that the species responded differently in the control versus dicamba environment. For the approach and visitation rates, contrasts were performed as indicated above, whereas for the binomial model assessing presence/absence of approaches or visitations, the risk ratios (ratios of probabilities) were determined in emmeans (64).

We controlled for shared evolutionary histories (65) by creating a phylogenetic tree (Fig S4) and used it to examine phylogenetically controlled models for each plant trait and the insect approach and visitation rates (see Methods S1 for details of analysis). However, models which controlled for phylogeny performed worse or similar compared to models that did not control for phylogeny based on Akaike information criterion (AIC) model selection (AIC values were typically 4 units higher; Table S4, (66)). We thus performed further analyses (structural equation modeling, below) and present results hereon from the non-phylogenetically corrected models only.

Finally, we performed path analysis using structural equation modeling (67) to examine hypothesized causal paths by which dicamba affects flower visitors. More specifically, we performed structural equation modeling to 1) examine the potential that dicamba exposure indirectly alters floral visitation rates by directly influencing plant traits (vegetative size, the presence of damage, day of first flowering, and flower display) and to 2) determine if interrelationships between plant traits are altered by the presence of dicamba drift. Structural equation modeling is a useful way to test direct and indirect effects by linking multiple variables into a single framework as well as a method that allows for simultaneous testing of multiple hypotheses (67, 68). We performed our analyses across weed species (69) and first developed an overall model to test treatment effects and then examined trait interrelationships within each treatment separately using the same general model structure (Fig S3).

To examine direct versus indirect effects of dicamba, we developed three hypothesized models (Fig S4). In the first model (Model A) we hypothesized that dicamba drift directly affects each of the measured plant traits – the amount of damage a plant exhibits, plant growth, flowering date, and total number of flowers – which leads to indirect effects on insect approach and visitation rates. Although we applied dicamba prior to plant flowering, and thus might not expect dicamba drift to directly impact flowering phenology or production, dicamba residues have been recovered in watermelon fruits and sweet potato storage roots when applied at drift rates (70, 71), suggesting that residues of this synthetic auxin could remain in plant reproductive tissues long after initial application or that plant hormonal pathways are altered long after application, also see (38). Our next hypothesized model (Model B) removed the potential for direct effects on flowering time and flower number such that any effects of dicamba exposure on both flowering time and flower number would be indirect and *via* direct effects on vegetative tissues. We included the effects of both plant damage and growth on flowering time and flower number in Models A and B, and likewise included the potential that flowering time influenced floral display, since plants that flower early typically produce more flowers over the season. In our final hypothesized model (Model C), we reduced Model B by removing the effect of growth on day of flowering and plant damage on total number of flowers since preliminary analyses found these relationships to be negligible, we had no *a priori* biological hypotheses regarding these relationships that would argue for their inclusion, and, using a model with fewer estimated parameters allowed us to model trait responses of plants from each treatment separately to examine how relationships between traits were altered due to dicamba drift.

We used piecewiseSEM (68) to apply structural equation modeling. The overall fit of each of the above hypothesized models was examined with Shipley’s d-generalized separation test, which is based on Fisher’s C statistics and evaluates if the data fit the hierarchical predictions proposed in the model and if other possible explanatory variables were excluded (72). We also report chi-square tests of model fit and select the most appropriate hypothesized model using Akaike information criterion (AIC) model selection. After we identified the most appropriate model (Model C, see results), we determined if interrelationships between traits were altered by the presence of dicamba by examining treatment-specific models, which follow the same general pattern of the overall model (Model C). We report model fit statistics for the SEMs performed in each treatment in SI Appendix.

---

**Table S1.** Collection site information for seeds used in this study.

| Site | Species | Latitude | Longitude | County, State in USA |
| --- | --- | --- | --- | --- |
| 15 | *Amaranthus palmeri* | 37.1281 | -86.6596 | Butler County, KY |
| 22 | *Amaranthus palmeri*, *Lepidium virginicum* | 36.7271 | -88.1781 | Calloway County, KY |
| 24 | *Taraxacum officinale*, *Daucus carota*, *Oxalis stricta* | 36.6614 | -88.2168 | Calloway County, KY |
| 20 | *Plantago virginica* | 36.8494 | -87.3016 | Christian County, KY |
| 21 | *Sida spinosa* | 36.854 | -87.5514 | Christian County, KY |
| 29A | *Plantago lanceolata* | 36.7995 | -87.3839 | Christian County, KY |
| 30A | *Oxalis stricta* | 36.7929 | -87.3796 | Christian County, KY |
| 27A | *Ipomoea hederacea* | 36.8257 | -87.477 | Christian County, KY |
| 17 | *Cardiospermum halicacabum* | 36.9771 | -86.7835 | Logan County, KY |
| 31 | *Sida spinosa*, *Abutilon theophrasti*, *Cardiospermum halicacabum* | 36.7014 | -87.0415 | Logan County, KY |
| 32A | *Oxalis stricta* | 36.6409 | -87.1576 | Montgomery County, TN |
| 11 | *Ipomoea lacunosa* | 36.5976 | -86.8511 | Robertson County, TN |
| 33A | *Plantago lanceolata*, *Daucus carota* | 36.44 | -86.8106 | Robertson County, TN |
| DG07 | *Amaranthus palmeri*, *Ipomoea lacunosa* | 36.6233 | -86.8413 | Robertson County, TN |
| DG08 | *Ipomoea hederacea* | 36.6319 | -86.8636 | Robertson County, TN |
| HJ01 | *Sida spinosa*, *Taraxacum officinale* | 35.751 | -86.6208 | Rutherford County, TN |
| HJ03 | *Ipomoea lacunosa*, *Cardiospermum halicacabum* | 35.7333 | -86.5902 | Rutherford County, TN |
| 19 | *Taraxacum officinale* | 36.8153 | -87.1768 | Todd County, KY |
| 14 | *Lepidium virginicum*, *Persicaria pennsylvanica* | 37.0655 | -86.6151 | Warren County, KY |

**Table S2.** Results from twelve rounds of recording insect visitor abundance and visitation data. In (A), results from an analysis of variance assessing insect abundance screens are presented whereas (B) shows the results of an analysis assessing approach and visitation rate (number of approaches or visitations/flower/hour) and (C) shows the results from analyzing approaches and visitations across observation rounds as a binary variable. In A, Wald Chisq values were determined using the ‘drop1’ function in R, with AIC estimates of model fit from dropping each explanatory variable presented. In B and C, type 3 Wald Chisq tests were performed using the anova function in car (57).

| A. | Abundance |  |  |  |
| --- | --- | --- | --- | --- |
|  | npars | AIC | Chisq | P |
| Treatment | 1 | 278 | 4.98 | **0.025** |
| Pollinator type | 9 | 382.48 | 125.45 | **<0.001** |
| Flower number | 1 | 276.47 | 3.44 | 0.06 |

| B. | Approach rate |  |  | Visitation rate |  |  |
| --- | --- | --- | --- | --- | --- | --- |
| *Fixed effects* | df | Chisq | P | df | Chisq | P |
| Treatment | 1 | 0.02 | 0.878 | 1 | 1.52 | 0.218 |
| Species | 7 | 28.58 | **<0.001** | 7 | 120.94 | **<0.001** |
| Treatment*Species | 7 | 4.34 | 0.739 | 7 | 21.07 | **0.004** |

| C. | Approach (y/n) |  |  | Visit (y/n) |  |  |
| --- | --- | --- | --- | --- | --- | --- |
| *Fixed effects* | df | Chisq | P | df | Chisq | P |
| Treatment | 1 | 0.00 | 0.95 | 1 | 1.88 | 0.171 |
| Species | 7 | 48.97 | **<0.001** | 7 | 167.41 | **<0.001** |
| Treatment*Species | 7 | 20.46 | **0.004** | 7 | 25.11 | **<0.001** |
| Flower Number | 1 | 6.54 | **0.011** | 1 | 5.52 | **0.019** |
| Plot | 1 | 5.08 | **0.024** | 2 | 7.13 | **0.03** |

**Table S3.**
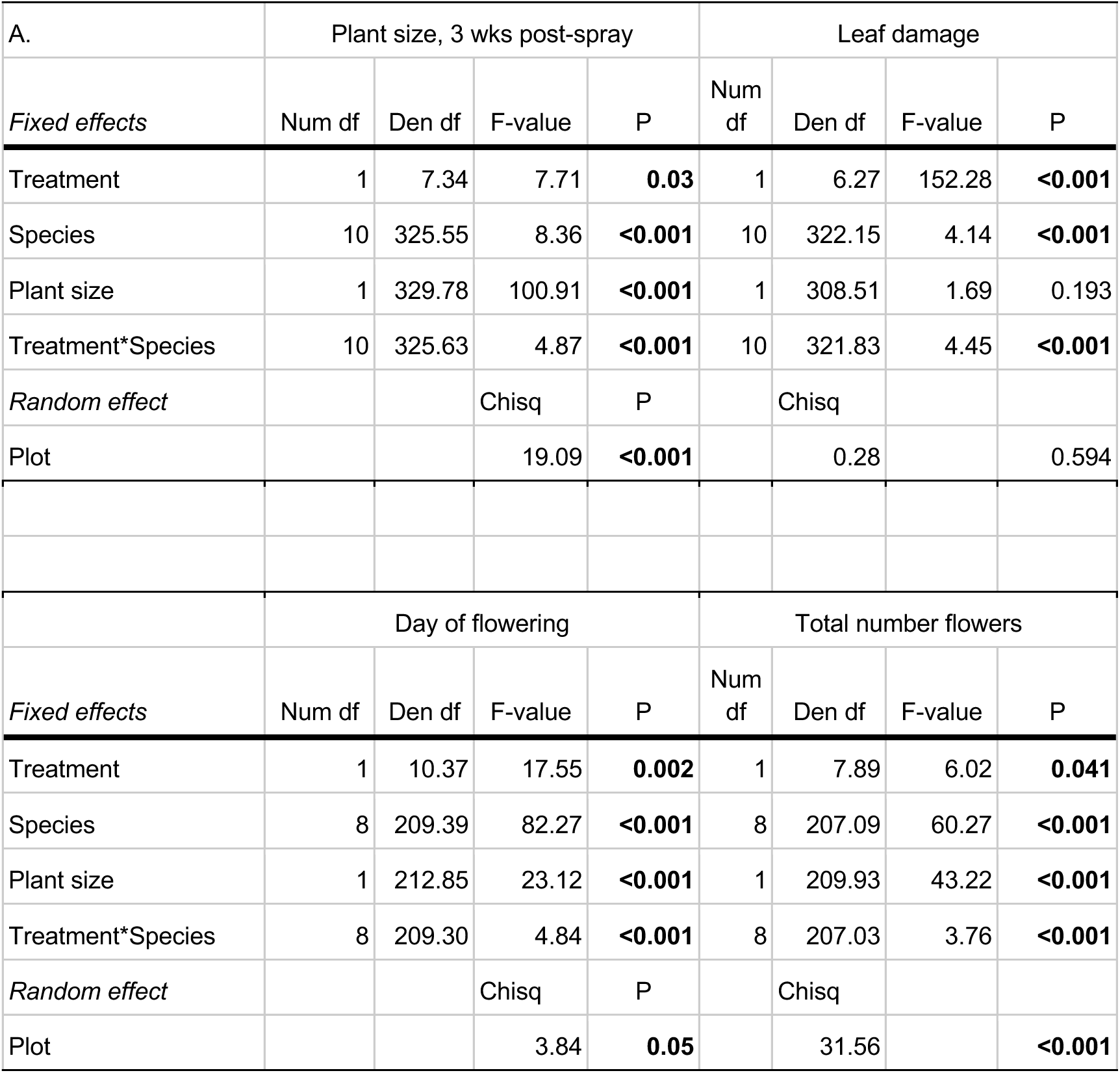
ANOVA type III tests for all measured plant traits. LmerTest (75) was used to assess significance of fixed effects with Satterthwaite’s method. The Wald chi-square statistic is reported in the ‘Chisq’ column for tests of random effects. The fixed effect ‘Treatment’ includes two levels (control and drift) and ‘Species’ includes 9-11 species, depending on the response (see Statistical Analysis in main text). Significant *P*-values are in bold.

| A. | Plant size, 3 wks post-spray |  |  |  | Leaf damage |  |  |  |
| --- | --- | --- | --- | --- | --- | --- | --- | --- |
| *Fixed effects* | Num df | Den df | F-value | P | Num df | Den df | F-value | P |
| Treatment | 1 | 7.34 | 7.71 | **0.03** | 1 | 6.27 | 152.28 | **<0.001** |
| Species | 10 | 325.55 | 8.36 | **<0.001** | 10 | 322.15 | 4.14 | **<0.001** |
| Plant size | 1 | 329.78 | 100.91 | **<0.001** | 1 | 308.51 | 1.69 | 0.193 |
| Treatment*Species | 10 | 325.63 | 4.87 | **<0.001** | 10 | 321.83 | 4.45 | **<0.001** |
| *Random effect* |  |  | Chisq | P |  | Chisq |  |  |
| Plot |  |  | 19.09 | **<0.001** |  | 0.28 |  | 0.594 |
|  | Day of flowering |  |  |  | Total number flowers |  |  |  |
| *Fixed effects* | Num df | Den df | F-value | P | Num df | Den df | F-value | P |
| Treatment | 1 | 10.37 | 17.55 | **0.002** | 1 | 7.89 | 6.02 | **0.041** |
| Species | 8 | 209.39 | 82.27 | **<0.001** | 8 | 207.09 | 60.27 | **<0.001** |
| Plant size | 1 | 212.85 | 23.12 | **<0.001** | 1 | 209.93 | 43.22 | **<0.001** |
| Treatment*Species | 8 | 209.30 | 4.84 | **<0.001** | 8 | 207.03 | 3.76 | **<0.001** |
| *Random effect* |  |  | Chisq | P |  | Chisq |  |  |
| Plot |  |  | 3.84 | **0.05** |  | 31.56 |  | **<0.001** |

**Fig S1.**
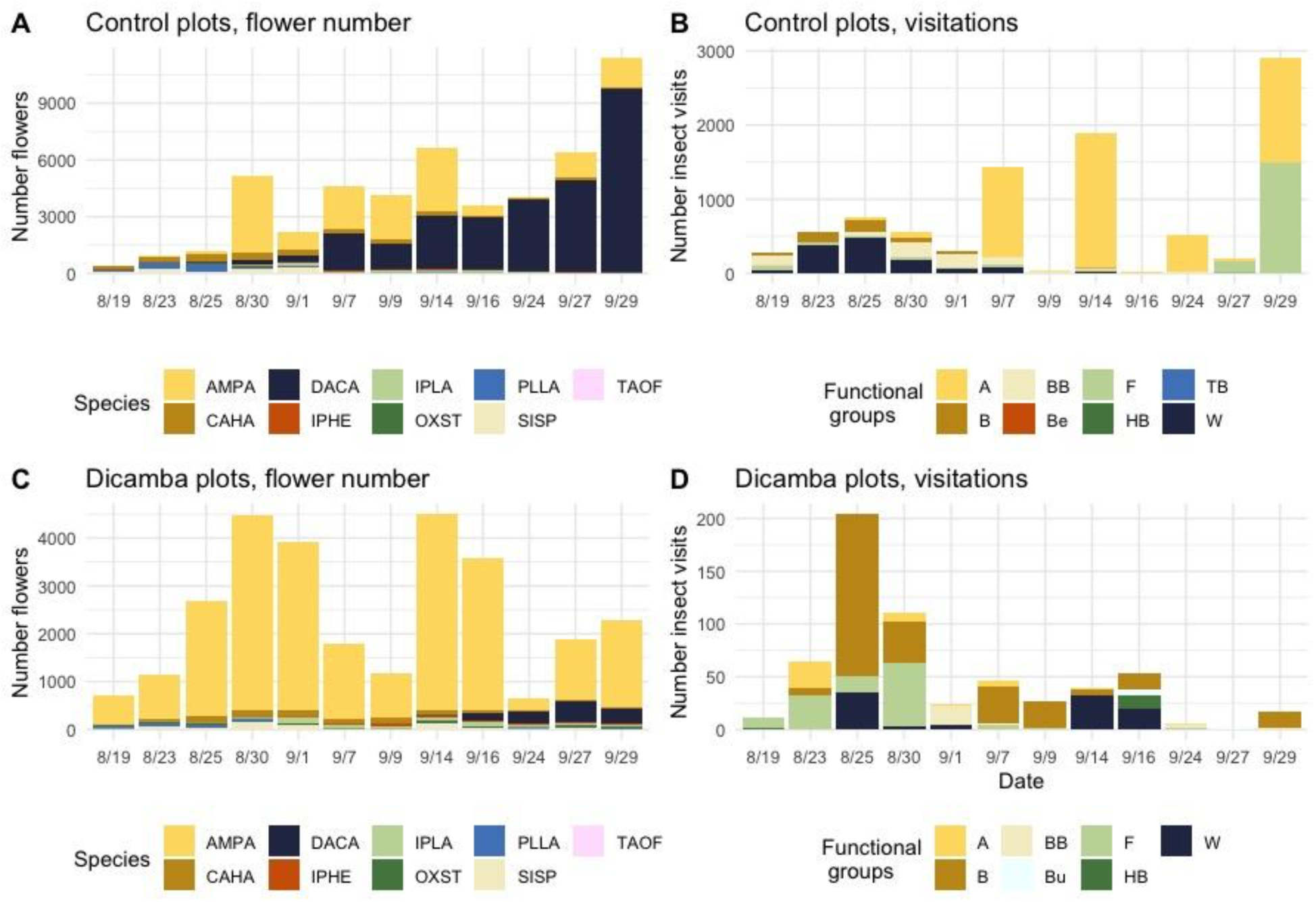
The number of flowers and insect visitations (forages within a flower) recorded over the course of the field experiment. The total number of flowers produced per species and the total number of visitations recorded per insect functional group type (Ant = ant, Fly = fly, Bee = Bee, TB = true bug, Wasp = wasp, BB = bumblebee, HB = honeybee, Be = beetle, Bu = butterfly)) in the control environment are shown in A and B, respectively, whereas C and D shows the same but in the dicamba treated plots.

**Fig S2.**
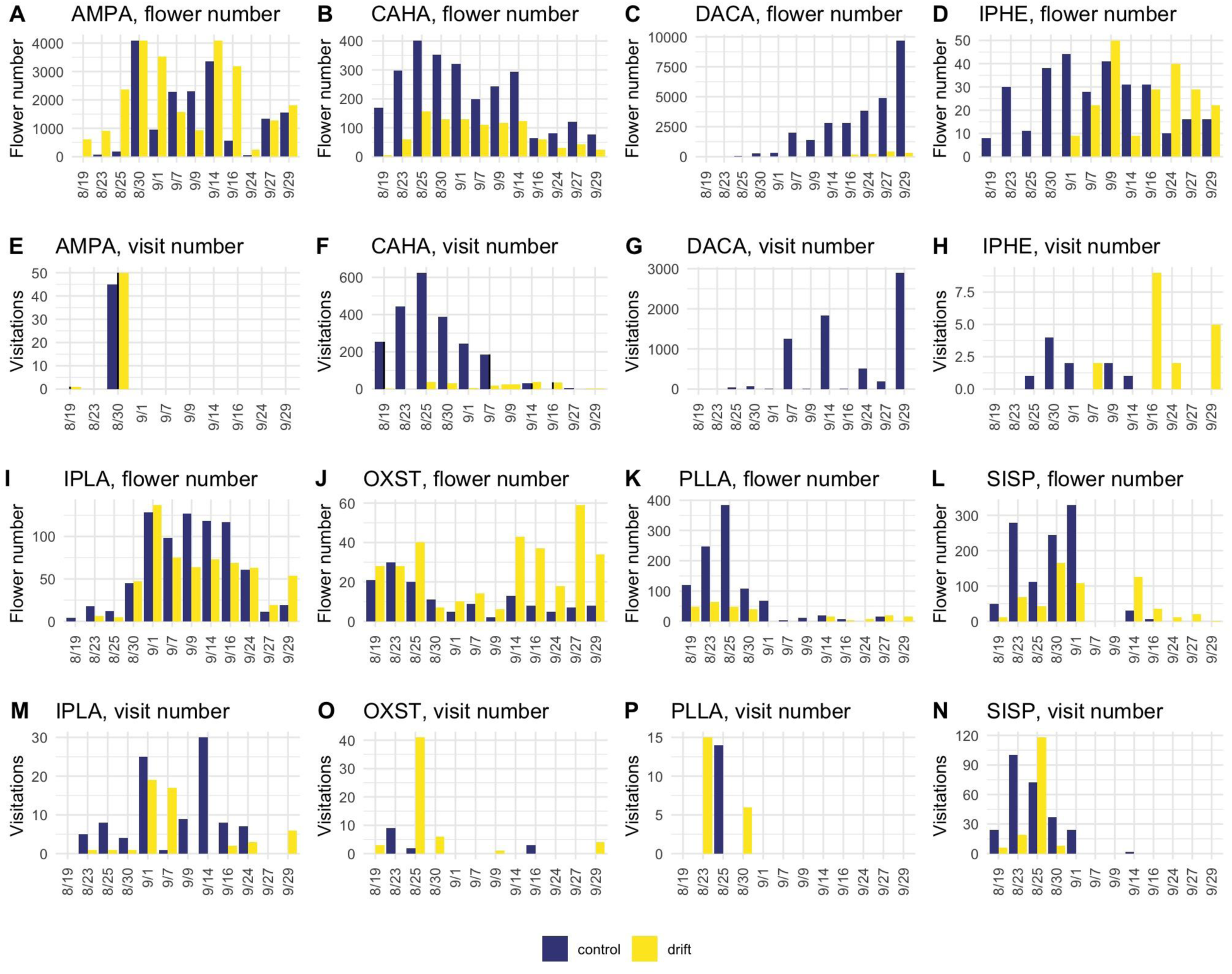
The total number of insect visits recorded across each visitation round and the total number of flowers produced during the experiment for each plant species. Summed visits and flowers of plants in control and dicamba exposed plots are shown in blue and yellow respectively. AMPA = *Amaranthus palmeri*; CAHA = *Cardiospermum halicacabum*; DACA = *Daucus carota*; IPHE = *Ipomoea hederacea*; IPLA = *Ipomoea lacunosa*; OXST = *Oxalis stricta*; PLLA = *Plantago lanceolata*; SISP = *Sida spinosa*; *Solidago canadensis* (SOSP) and *Taraxacum officinale* (TAOF) were removed from analyses since fewer than two plants produced flowers in the presence of dicamba drift.

Methods S1. We controlled for shared evolutionary histories (65) by creating a phylogenetic tree (SFig 3) with the phytools (76) and V. PhyloMaker (77) packages. We extracted phylogenetic information from the mega-phylogenetic tree ‘GBOTB’ for seed plants (78) and, for all response variables, used this tree to perform tests of phylogenetic models (Table S4) with the phylolm function (phylolm package; (79) and the pglmm_compare function (phyr package; (80)). Using the reconstructed phylogenetic trees, we determined if phylogenetically-corrected mixed models exhibited lower AIC values,and thus were the best-fit, compared to the non-phylogenetically corrected models (66).

**Fig S3.**
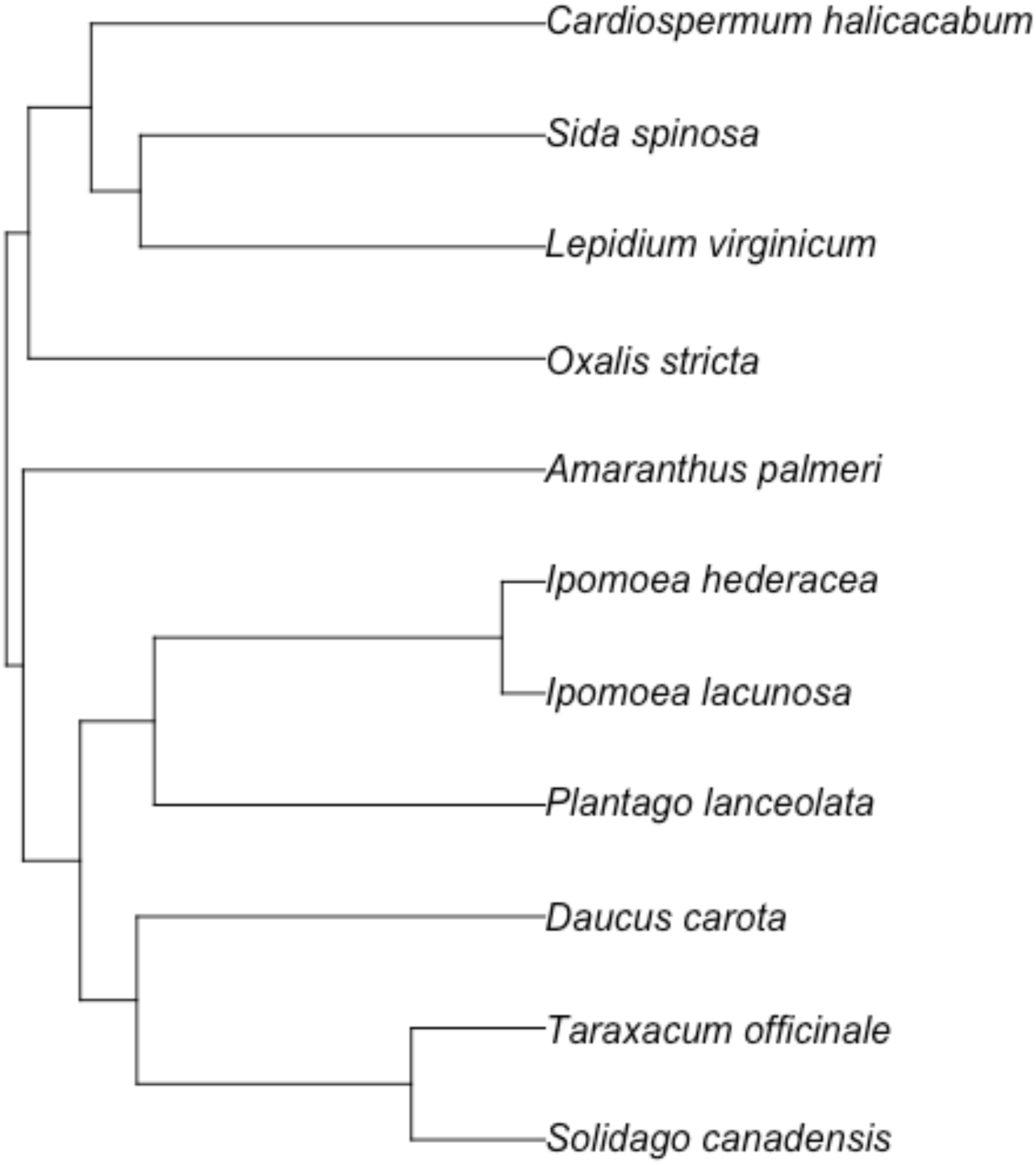
Phylogeny of the 11 plant species used in this study. We constructed the tree using the R package V.PhyloMaker (77). We used the options tree = GBOTB.extended (indicates that phylogenetic information was extracted from a corrected combination of the mega-phylogenetic tree ‘GBOTB’ for seed plants (78)), nodes = nodes.info1 (genus- and family-level node and branch length information provided by the mega-tree), and scenarios = “S1” (new species tips are binded to genus- or family-level basal nodes).

**Table S4.** Estimates of goodness of fit for each model in this study based on Akaike information criterion (AIC) with and without phylogenetic correction.

|  | Phylogenetic model |  | Non-phylogenetic model |  |  |
| --- | --- | --- | --- | --- | --- |
| Model | AIC | logLik | AIC | logLik | P |
| asinh(Size 3 wks post treatment) ~ Treatment x Species + Pretreatment size + (1\|treatment/plot) | 1033.2 | -489.6 | 1029.2 | 497.3 | 1 |
| sqrt(1+Leafdamage) ~ Treatment x Species + Pretreatment size + (1\|treatment/plot) | -940.5 | 497.3 | -944.5 | 497.3 | 0.987 |
| sqrt(Day of first flower) ~ Treatment x Species + Pretreatment size + (1\|treatment/plot) | 163.37 | -58.68 | 159.37 | -58.68 | 1 |
| asinh(Number flowers) ~ Treatment x Species + Pretreatment size + (1\|treatment/plot) | 678.9 | -316.5 | 674.9 | -316.5 | 0.963 |
| (1+Visit Rate) ~ Treatment x Species + (1\|treatment/plot) | -130.45 | 87.22 | -134.45 | 87.22 | 0.993 |
| (1+Approach Rate) ~ Treatment x Species + (1\|treatment/plot) | -260.1 | 152.1 | -264.1 | 152.1 | 0.993 |

**Fig S4.**
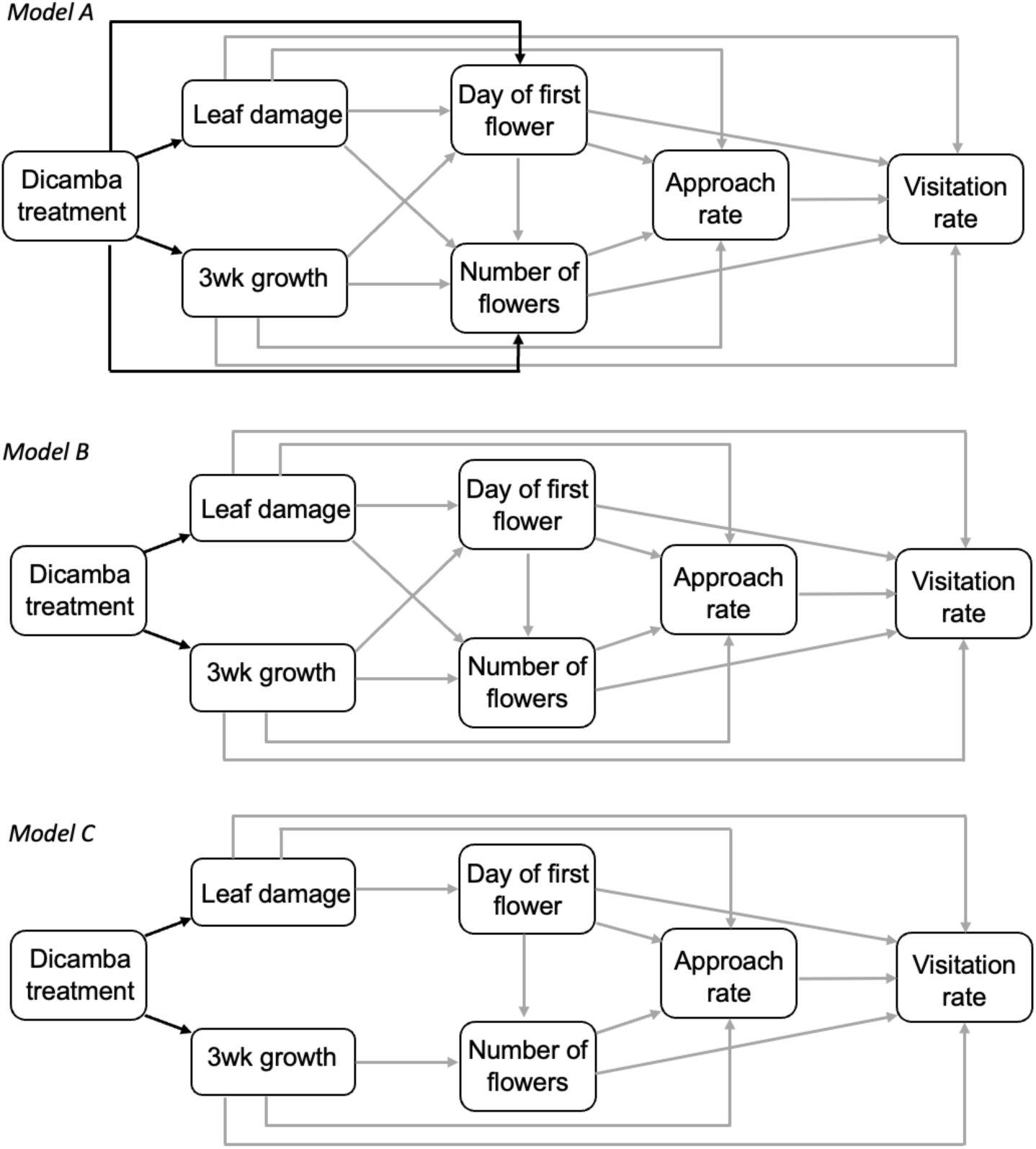
Three hypothesized structural equation models examined in this work. The full model (Model A) examined the hypothesis that dicamba exposure (dicamba treatment) indirectly affects visitation rate through direct effects on each of the plant traits (leaf damage, size at 3 weeks (3 wk growth), number of flowers, and day of first flower). In Model B, we hypothesized that dicamba exposure significantly and directly influences plant vegetative traits (leaf damage, 3 wk growth) but all other effects are indirect. In Model C, we further reduced model B by removing the pathways between leaf damage and number of flowers, and 3 wk growth and day of first flower. Preliminary examination suggested the models were not significantly different from one another, and effects were not significant.

**Table S5.** SEM model comparisons. piecewiseSEM (68) was used to perform model fit tests (Chisq, Fisher’s C) and contrast models. Each hypothesized model fit the data, and none of the models (A-C) varied significantly from one another (p > 0.05 each contrast). The AIC value of each model is presented as are the number of parameters (k) and sample size for each model (n).

| Model | Description | AIC | k | n | Chisq | P | FisherC | df | P |
| --- | --- | --- | --- | --- | --- | --- | --- | --- | --- |
| A | Full model | 1266.59 | 30 | 207 | 0.6 | 0.9 | 2.48 | 6 | 0.87 |
| B | No direct effect of dicamba on flowering day or flower number | 1264.5 | 28 | 207 | 2.53 | 0.77 | 6.52 | 10 | 0.77 |
| C | B, and no effect of leaf damage or growth on flower number or day of flowering, respectively | 1261.32 | 26 | 207 | 3.35 | 0.85 | 9.01 | 14 | 0.83 |
| control only | model C within control | 512.98 | 20 | 101 | 3 | 0.22 | 5.86 | 4 | 0.21 |
| dicamba only | model C within dicamba | 416.91 | 20 | 106 | 1.65 | 0.44 | 3.17 | 4 | 0.53 |

**Table S6.** The pathways leading to indirect effects of dicamba exposure on pollinator visit rate. For each pathway, the standardized path coefficients were multiplied, and total effects were determined by summing across pathways.

| Pathway | Description of pathway | Total effect |
| --- | --- | --- |
| 1 | Dicamba treatment→leaf damage→day of flowering→approach rate<br>$0.62 \times 0.27 \times -0.27 \times 0.40$ | -0.02 |
| 2 | Dicamba treatment→3wk growth→flower number<br>$-0.38 \times 0.61 \times 0.27$ | -0.06 |
| 3 | Dicamba treatment→3wk growth→approach rate<br>$-0.38 \times 0.57 \times 0.40$ | -0.09 |
|  | Grand Total effect | -0.17 |

